# ADC mapping with 12 *b* values: an improved technique for image quality in diffusion prostate MRI

**DOI:** 10.1101/744961

**Authors:** Lucas Scatigno Saad, George de Queiroz Rosas, Homero José de Farias e Melo, Henrique Armando Azevedo Gabriele, Jacob Szejnfeld

**Affiliations:** Universidade Federal de São Paulo (UNIFESP), São Paulo, SP, Brazil

## Abstract

**Purpose:** To compare diffusion images and coefficients obtained with 4 *b*-value versus 12 *b*-value apparent diffusion coefficient (ADC) mapping for characterization of prostate lesions and how these coefficients relate and compare to the PI-RADS™ classification and Gleason grading system.

**Methods:** Patients with indications for prostate cancer testing (n=158) underwent multiparametric 3T magnetic resonance imaging (MRI). Two diffusion sequences were acquired, one with 4 *b* values and one with 12 *b* values. ADC maps were calculated for each (ADC_4_ and ADC_12_) and the respective coefficients were tested for correlation with PI-RADS™ classification and Gleason score.

**Results:** The ADC_12_ sequence produced images of superior quality and sharpness than ADC_4_. Normal-area means (ADC_4_, 1793.3×10^−6^mm^2^/s; ADC_12_, 1100×10^−6^mm^2^/s) were significantly lower than those of lesion areas (ADC_4_, 1105.9×10^−6^mm^2^/s; ADC_12_, 689.4×10^−6^mm^2^/s) (p<0.001). Both techniques behaved similarly and correlated well with PI-RADS™ classification, distinguishing scores 3, 4, and 5 and with means tending to decline with increasing Gleason grade. ADC_12_ mapping yielded higher specificity than ADC_4_ (82.6% vs. 72.3%).

**Conclusions:** Diffusion with 12 values is a viable technique for examination of the prostate. It produced higher-quality images than current techniques and correlates well with PI-RADS™ classification and Gleason score.

## Introduction

Prostate cancer is the third most prevalent cancer in the United States. About 160,000 new cases were diagnosed in 2017, representing approximately 10% of all new cases of cancer in men [1]. Classically, screening for prostate in the general population is performed by PSA (prostate specific antigen) testing and rectal examination; however, both are unsatisfactory for early detection of clinically significant lesions [2, 3].

Magnetic resonance imaging (MRI) for prostate evaluation was first proposed in the mid-1980s [4, 5], with the objective of staging already-diagnosed tumors; technological advances at the time enhanced the potential of MRI to detect suspicious lesions. Since then, MRI has gained widespread use in clinical practice, whether for screening or staging of tumors. MRI often avoids more invasive and unnecessary procedures, such as prostate biopsy, which is performed indiscriminately in most centers and can lead to a series of complications [6]. New techniques, such as biopsy with sonographic and resonance-derived fusion images, have become the focus of much recent literature due to their potential to increase diagnostic value [7, 8].

Among the tools used to diagnose prostate cancer by MRI, one stands out: diffusion-weighted imaging (DWI), a functional imaging sequence that measures signal arising from movement of water molecules in the tissue of interest. DWI requires at least two acquisitions with different time lengths and gradient amplitudes, commonly known as the *b* value, which is the measure of the diffusion power of a given sequence. Higher *b* values are associated with greater diffusion gradient weighting used in the sequence and, consequently, greater the diagnostic accuracy to study restriction. The effective limit is 2500 s/mm^2^[9], a threshold over which image distortion begins to occur with substantial loss of lesional spatial resolution.

The signal difference between diffusion sequences acquired with different *b* values results in a first-degree exponential equation that yields the apparent diffusion coefficient (ADC). The ADC, which is a quantitative measure of the movement speed and restriction of a given molecule, is expressed in square millimeters per unit time in seconds. It is a reproducible measure of diffusion that can be obtained at any dedicated workstation by measuring a delimited region of interest (ROI).

With the advent of the second version of PI-RADS™, an imaging classification that aims to stratify prostate imaging findings according to severity and risk [10, 11] for tumors of the peripheral zone, DWI has become the key sequence for approximating severity of a possible focal lesion.

In addition, the correlation between diffusion, as expressed as a numerical ADC value, and the aggressiveness of prostate tumors has also been widely studied in the literature. Significant diffusion restriction is associated with high histological aggressiveness, as measured by Gleason scores. Therefore, ADC correlates closely with prognosis and treatment planning for these patients [12–14].

However, ADC values obtained in diffusion sequences with *b* values up to 1000 s/mm^2^ show an overlap between normal tissues and neoplastic lesions [15–17]. There is no consensus in the current literature on what the discriminatory *b* value ought to be for standard study of the prostate [18, 19]. Recent studies show that high *b* values (up to 2000 s/mm^2^) are more sensitive for lesion identification. However, technique limitations affect the quality and sharpness of images obtained on ADC due to distortion [20–22].

This limitation inherent to DWI motivated the search for technical improvements to optimize image quality and resolution. One such improvement is the possibility of increasing acquired *b* values. In our experience, diffusion sequences performed with 12 *b* values provide much higher image quality than standard diffusion sequences, which are usually performed with 4 *b* values.

However, in order to include this sequence in routine prostate MRI examination, it must first be validated, especially with regard to technical parameters, quality, and diagnostic value of the obtained image.

Diffusion sequences are a key part of multiparameter (mp)-MRI study, and meet the appropriate criteria for diagnosis of prostate cancer. However, obtained images have technical and morphological limitations that often hinder proper identification and exact localization of lesions. Thus, seeking to improve the quality of anatomical visualization in diffusion sequences without sacrificing diagnostic capacity, a protocol using 12 *b* values were developed and compared it to the standard diffusion technique. The main objectives of this study are to evaluate the sharpness and conspicuity of images obtained in diffusion sequences with 4 and 12 *b* values for evaluation of the normal prostate and in the characterization of prostatic lesions; and to establish whether ADC measurements in both techniques correlate with prostate tumor aggressiveness.

## Materials and methods

### Sample

Multiparametric MRI of the prostate was performed in 158 patients with a clinical/laboratory indication for prostate cancer screening. Patients with increased PSA and/or altered rectal examination deemed clinically significant were included. Patients with known cancer who had a clinical indication for MRI staging were also included. Examinations were carried out between September 2015 and August 2016.

### Technical Parameters

Scans were performed on 3T equipment with a 45 mT/m gradient (Magnetom Verio and Magnetom Skyra; Siemens Medical Systems, Erlangen, Germany) using a standard torso coil.

The MRI sequences are summarized in Table 1. Axial spin-echo T2, coronal and sagittal T2 for morphological study of the prostate (256 × 230 matrix, 3.0 mm slice thickness, 160 × 160 mm FOV, TR = 3560 ms and TE = 114 ms), T1 axial spin-echo (256 × 230 matrix, 3.0 mm slice thickness, 160 × 160 mm FOV, RT = 550 ms and TE = 9.5 ms).

**Table 1.**
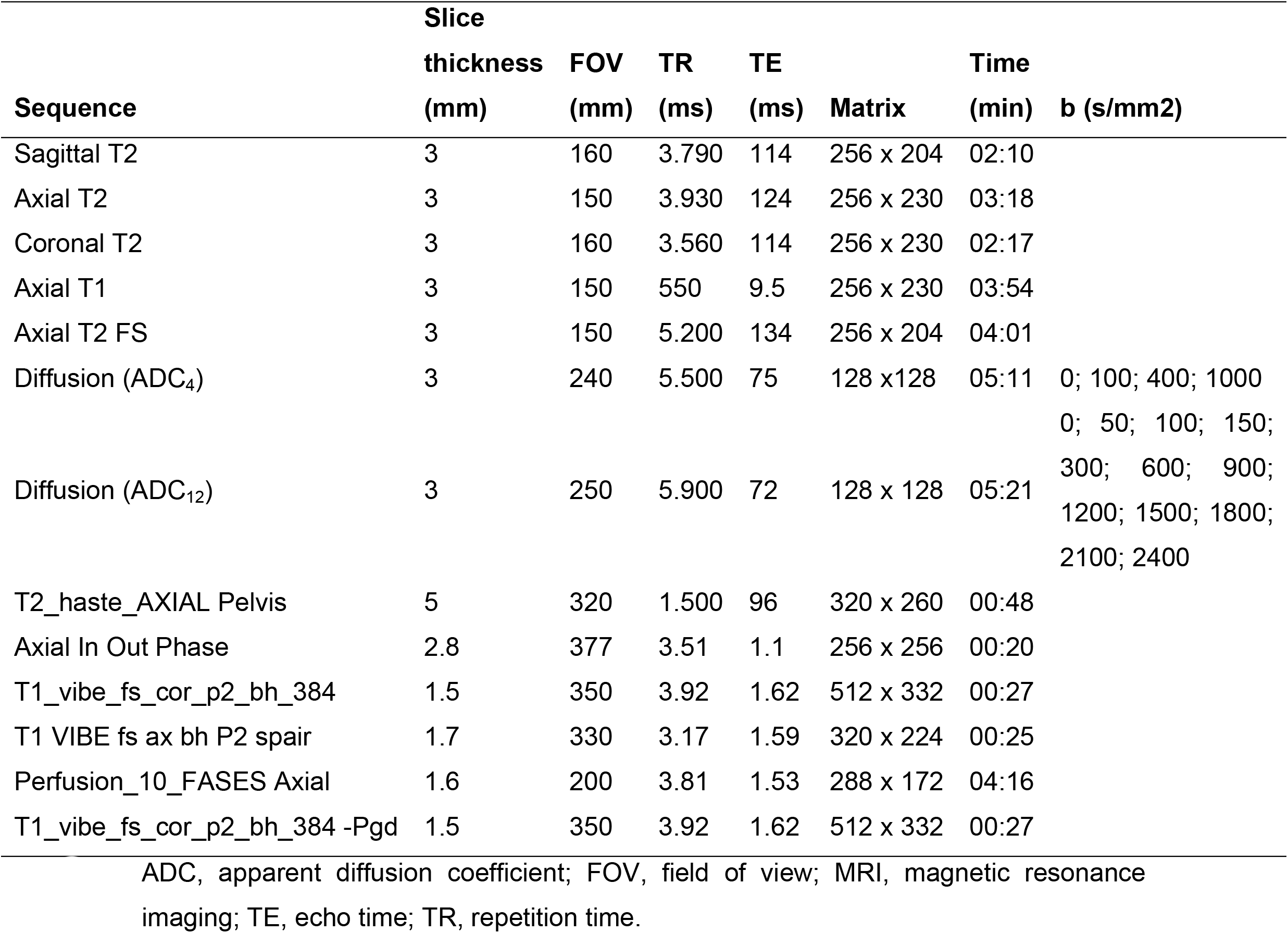
MRI sequences used in the study protocol.

Two single-shot echo-planar axial sequences with diffusion-weighted gradients for functional study of the prostate were employed: “diffusion 4” (128 × 128 matrix, 3.0 mm slice thickness, 240 × 240 mm FOV, TR = 5500 ms and TE = 75 ms with four values of *b* = 0; 100; 400; 1000 s/mm) and “diffusion 12” (128 × 128 matrix, 3.0 mm slice thickness, 240 × 240 mm FOV, TR = 5900 ms and TE = 72 ms with 12 values of *b* = 0, 50, 100, 150, 300, 600, 900, 1200, 1500, 1800, 2100, 2400 s/mm). Using acceleration tools, the diffusion 12 sequence lasted approximately 10 seconds longer than the diffusion 4 sequence (5 min 21 s versus 5 min 11 s) was obtained. In addition, when there was no absolute contraindication, the standard protocol included pre and post-contrast dynamic sequences.

Post-processing was then performed to calculate ADC maps with 4 values of *b* (ADC_4_) and with 12 values of *b* (ADC_12_).

### Imaging

Images from morphology and functional sequences were evaluated synchronously and simultaneously on a dedicated workstation (syngo.via, Siemens™) and analyzed in consensus between two radiologists with experience in prostate imaging. The following findings were considered as imaging criteria for a clinically significant prostatic lesion (suspicion for cancer): focal hypointensity in T2 and/or signal restriction in ADC_4_ and/or ADC_12_.

Measurements of ADC_4_ and ADC_12_ values were performed using the ROI tool in areas identified as suspicious, allowing for the largest possible lesion area and copying the same area to the ADC_4_ and ADC_12_ (Fig 1) through a specific tool that duplicated the ROI measurement for the sequence of interest. In the absence of a lesion, measurements were performed only on areas of normal prostate.

**Fig. 1.**
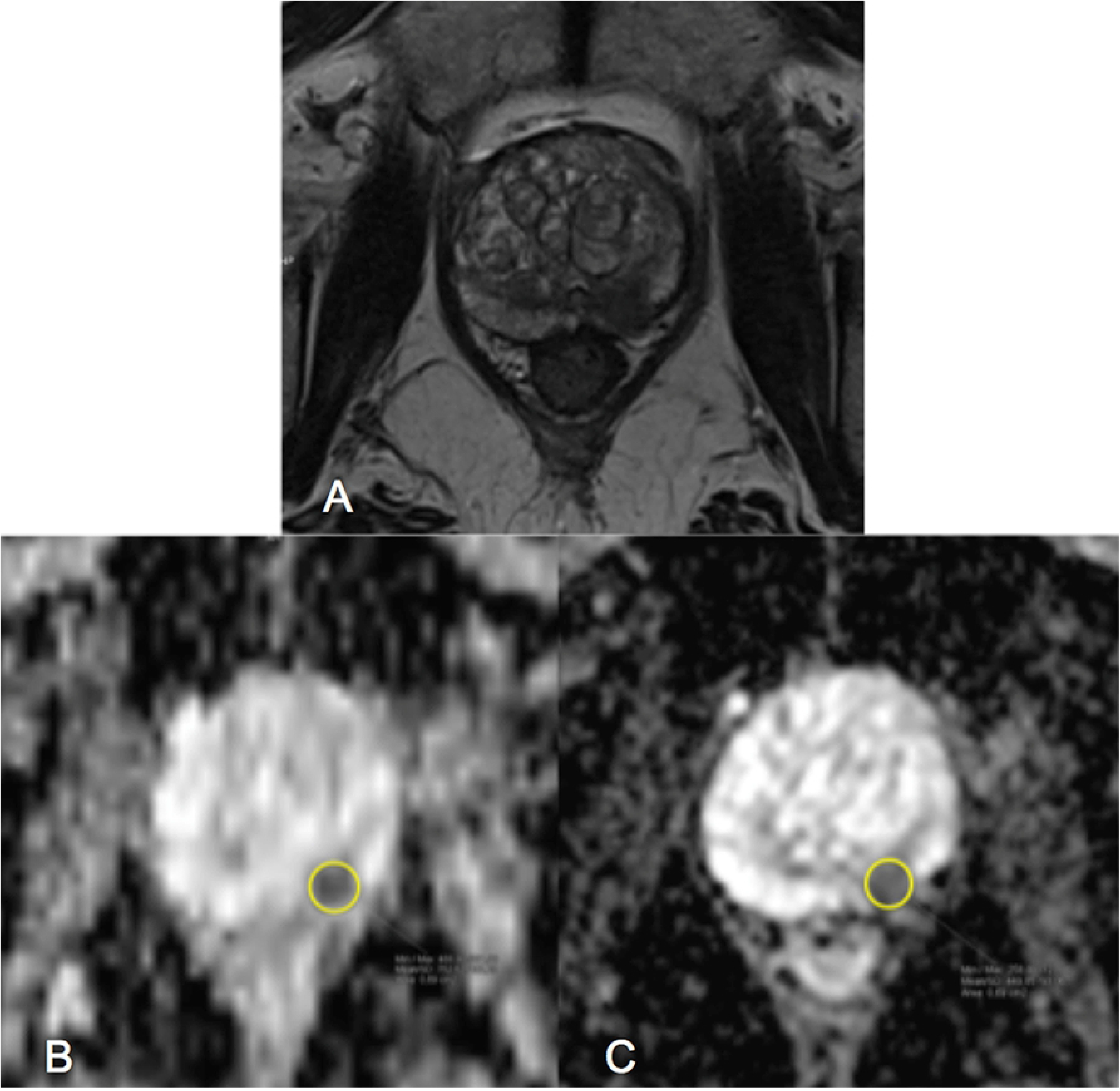
Representative magnetic resonance imaging in patient with a prostate mass. **A)** T2 sequence images showing morphological appearance of mass in left peripheral zone. **B)** ADC_4_ sequence; lesion exhibits restricted diffusion. Yellow circle represents ROI. **C)** ADC_12_ sequence; lesion exhibits restricted diffusion. Yellow circle represents ROI. ADC, apparent diffusion coefficient; ROI, region of interest.

To assess image quality, the two observers independently analyzed the sequences side by side on the workstation, grading the sharpness and conspicuity of two parameters – prostate anatomy and lesion visualization (when present) – on a scale of 1 (very low sharpness) to 5 (excellent sharpness).

### Statistical Analysis

The means, medians, and standard deviations of the ROI measurements of ADC_4_ and ADC_12_ were calculated for lesion areas and normal areas. Student’s *t*-test was used for comparison of signal behavior between normal and lesion areas. A regression model was used to compare measurements obtained in ADC_4_ and ADC_12_, as well as to test for correlation between average PI-RADS™ and Gleason classification when the patient underwent biopsy. Receiver operating characteristic (ROC) curves were used to calculate the sensitivity and specificity of the parameters of interest for prediction of cancer.

Likewise, the scores assigned to image quality were tabulated, analyzed, and compared between the two sequences also between the two observers.

## Results

According to the inclusion criteria, 51 patients presented with suspicious lesions that were measurable by the methodology used in the study design. Normal areas were measured both in patients with lesions and in patients without lesions, for a total of 158.

Means, medians, standard deviations and ranges are summarized in Table 2. Analysis of means and medians revealed higher values in the normal areas and smaller values in the lesion areas. In addition, absolute values for the ADC_12_ sequence were overall smaller compared to the ADC_4_ values, as observed in the minimum and maximum values for each measurement.

**Table 2.** ADC_4_ and ADC_12_ values*

| Parameters | Normal ADC<sub>4</sub> | Lesion ADC<sub>4</sub> | Normal ADC<sub>12</sub> | Lesion ADC<sub>12</sub> |
| --- | --- | --- | --- | --- |
| Mean | 1793.3 | 1105.9 | 1100.0 | 689.4 |
| 95% FI for mean |  |  |  |  |
| Lower limit | 1748.2 | 1022.0 | 1071.9 | 642.6 |
| Upper limit | 1838.3 | 1189.8 | 1128.1 | 736.1 |
| Median | 1829.5 | 1162.0 | 1092.5 | 680.0 |
| Standard deviation | 286.4 | 298.3 | 178.7 | 166.1 |
| Minimum | 1057.0 | 478.0 | 665.0 | 260.0 |
| Maximum | 2631.0 | 1695.0 | 1461.0 | 947.0 |

\* Values expressed as 10<sup>-6</sup> mm<sup>2</sup>/s.
ADC, apparent diffusion coefficient; FI, fiducial interval.

Comparison between mean ADC_4_ values in normal versus lesion areas revealed significantly higher coefficients in the former (Student’s *t* test for paired samples, p<0.001) (Fig 2). A similar relationship was also observed for values obtained from ADC_12_, with significantly higher means in normal versus lesion areas, thereby demonstrating similar behavior in the two techniques (Fig 3).

**Fig. 2.**
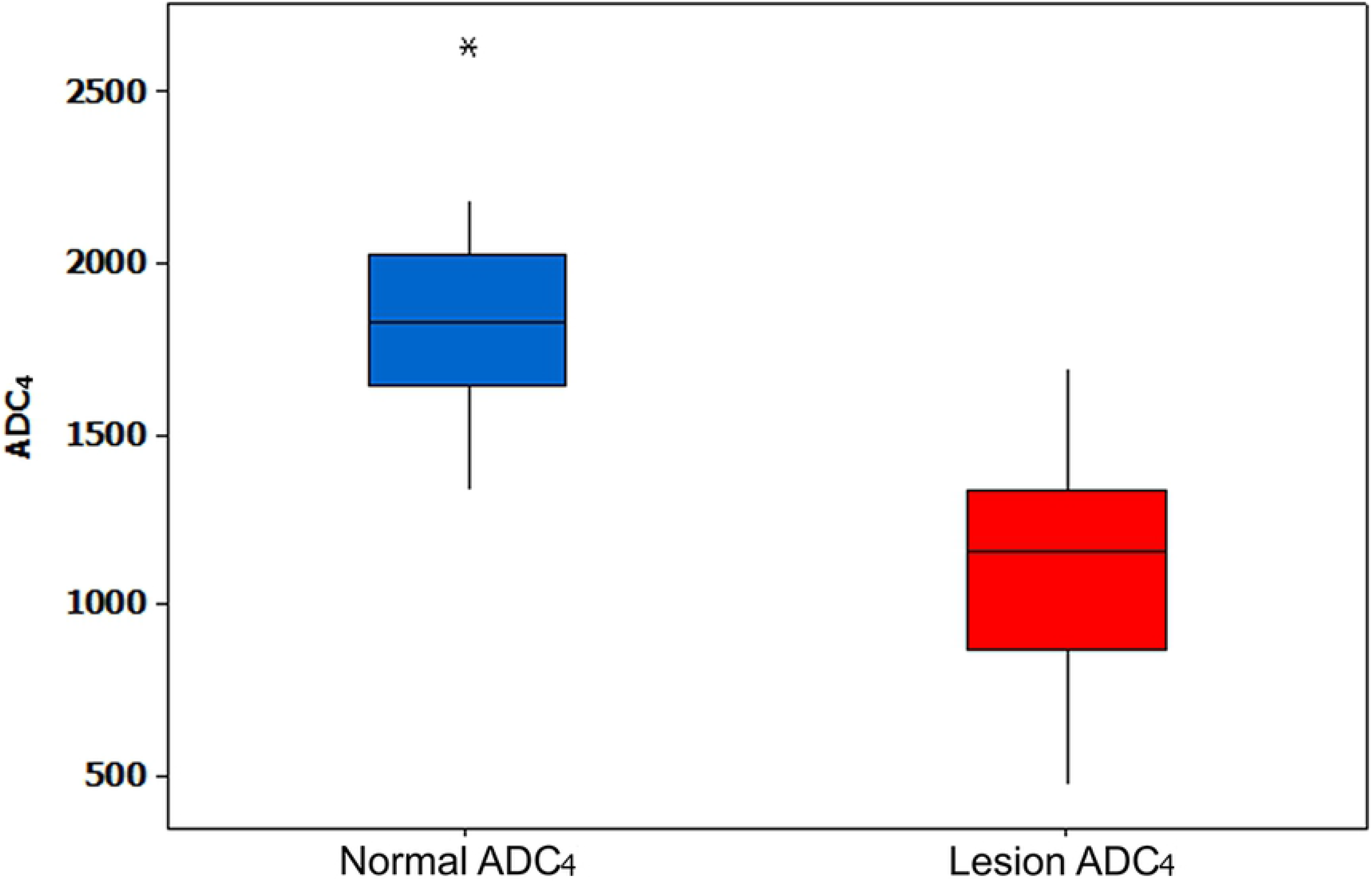
Box-plot of Normal ADC_4_ and Lesion ADC_4_. ADC, apparent diffusion coefficient.

**Fig. 3.**
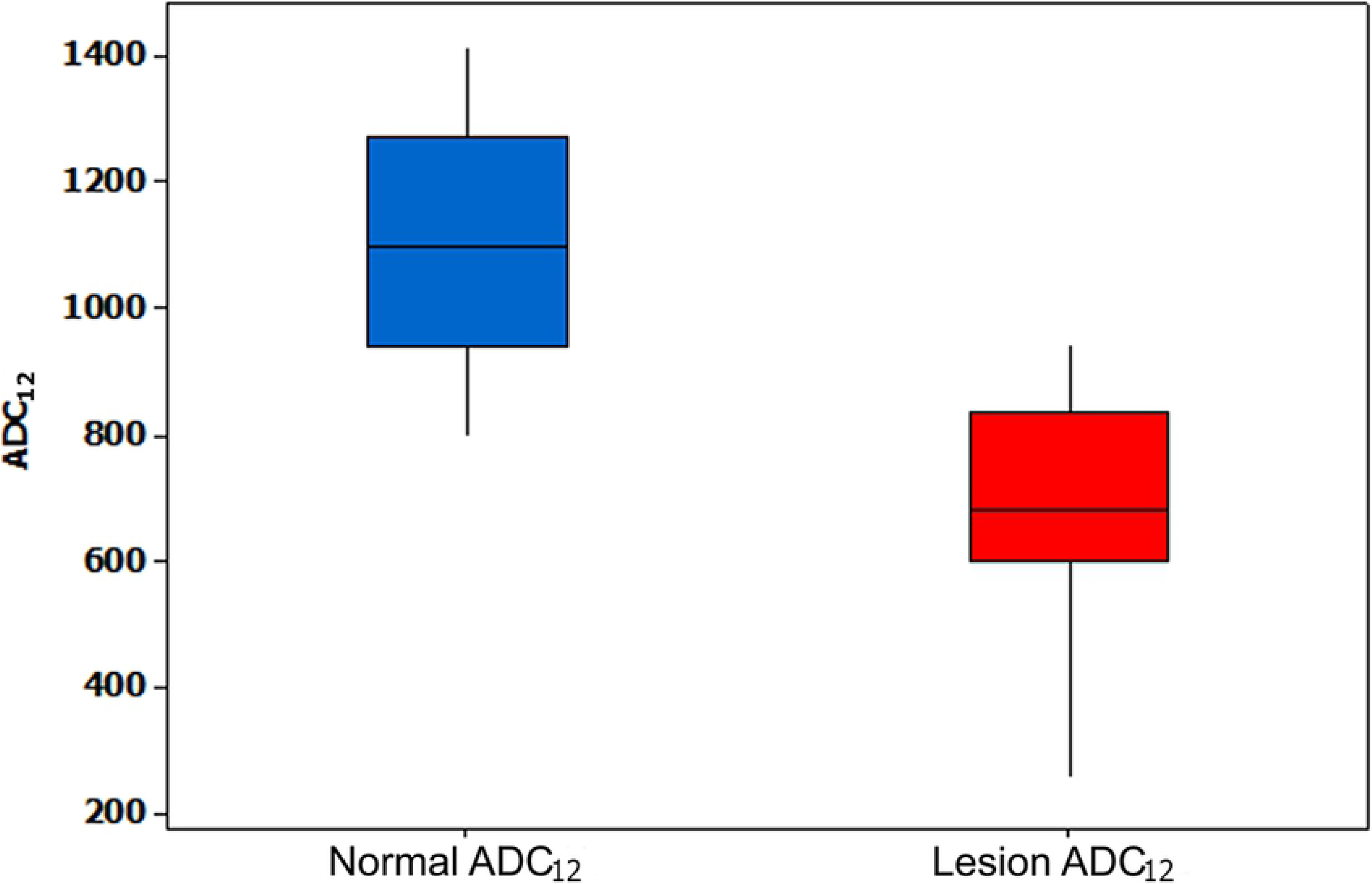
Box-plot of Normal ADC_12_ and Lesion ADC_12_. ADC, apparent diffusion coefficient.

Given this similarity in behavior between the two techniques, measurements were analyzed through a regression model between normal areas and lesion areas (Fig 4), which demonstrated a constant correlation in lesion measurements of the standard ADC_4_ and ADC_12_. The following mathematical regression formula was obtained:

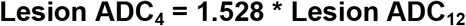

**Fig. 4.**
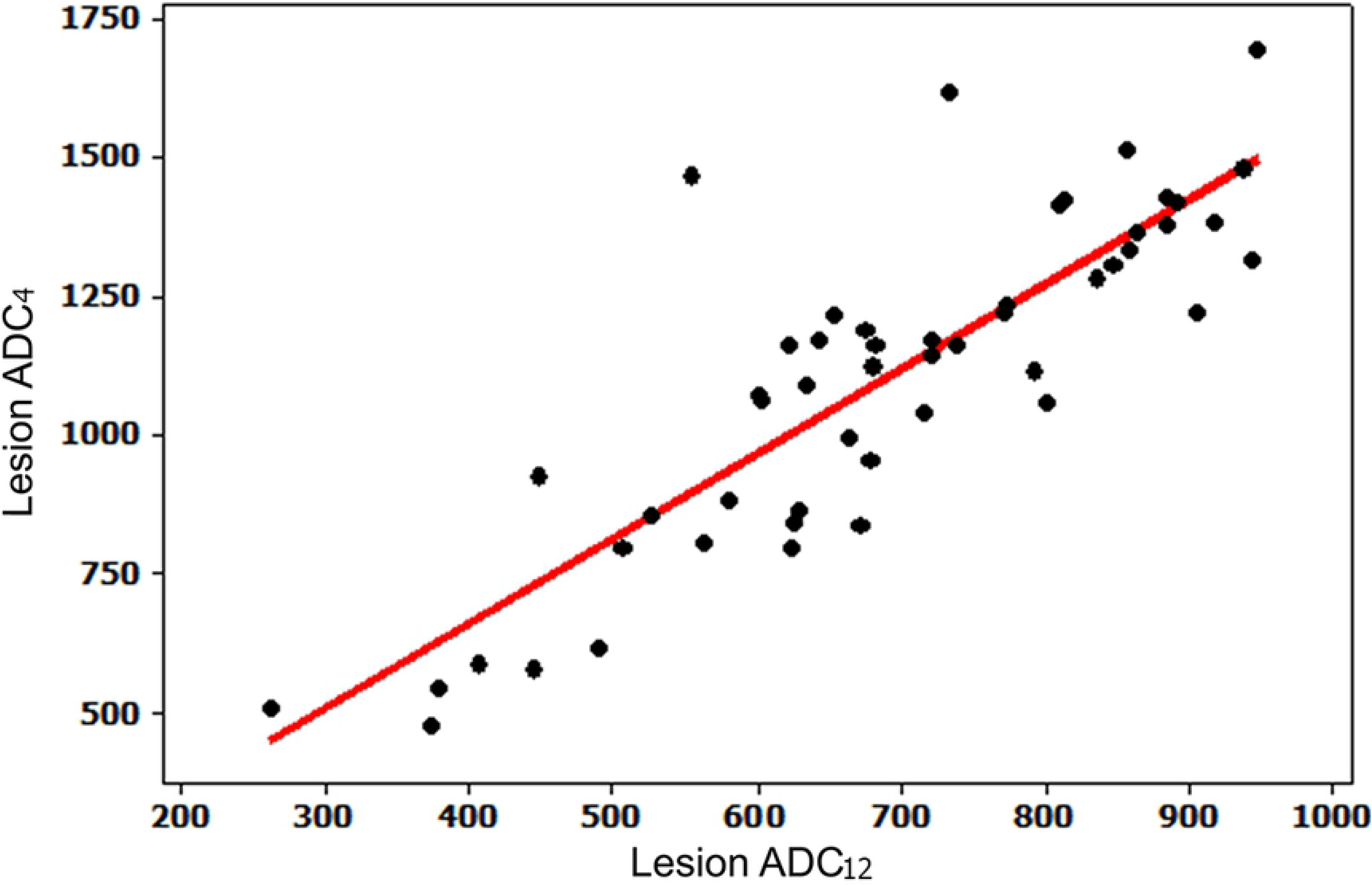
Scatterplot of Lesion ADC_4_ and Lesion ADC_12_. ADC, apparent diffusion coefficient

A similar correlation was also observed in the measurement of normal areas with standard ADC_4_ and ADC_12_ (Fig 5). By applying the same regression model, we obtained the following correlation formula:

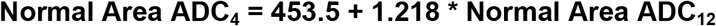

**Fig. 5.**
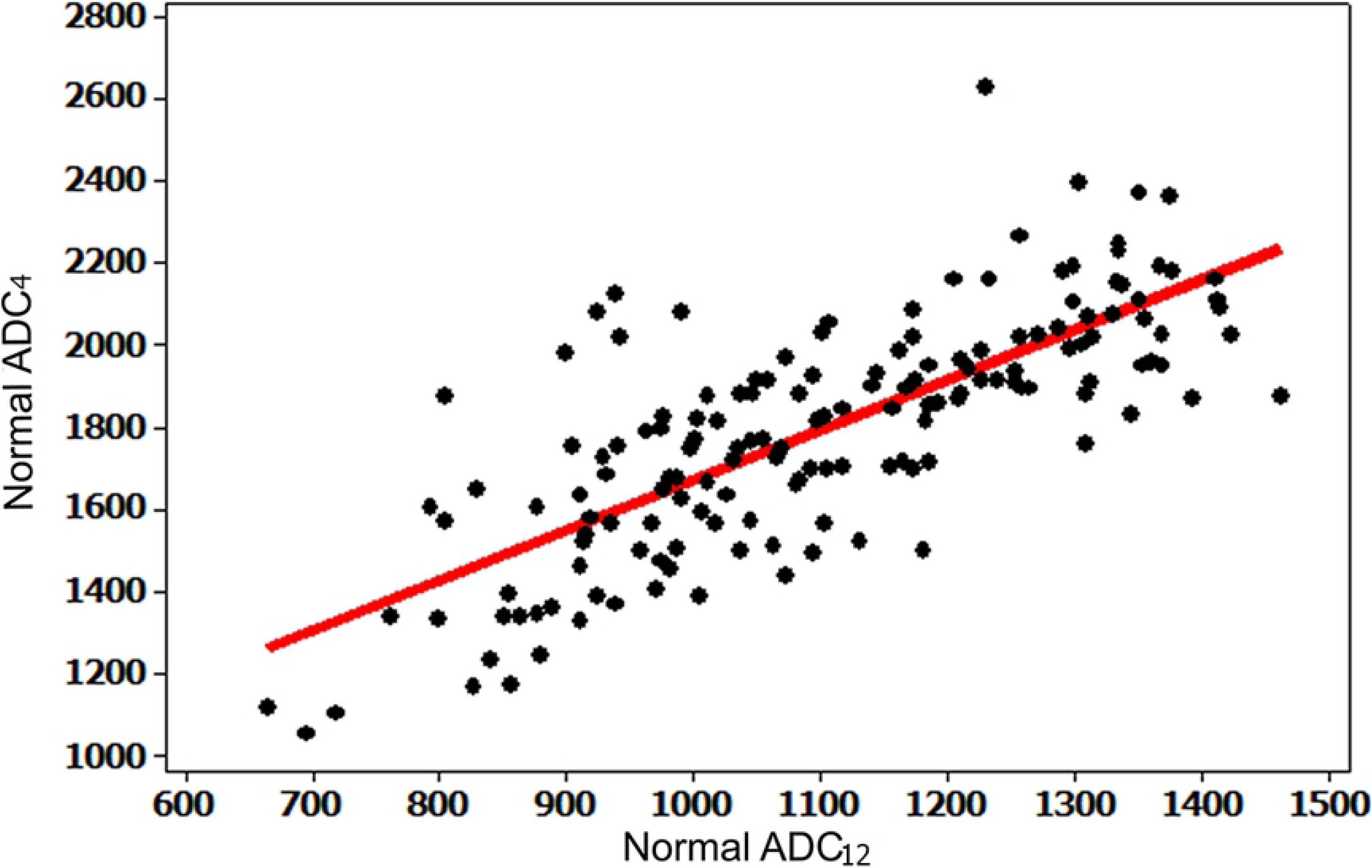
Scatterplot of Normal ADC_4_ and Normal ADC_12_. ADC, apparent diffusion coefficient

On PI-RADS™ v2 assessment, 6 patients were classified as category 1 (absence of clinically significant lesion), 99 as category 2 (low probability of clinically significant cancer), 11 as category 3 (presence of clinically significant cancer is equivocal), 36 as category 4 (high probability of clinically significant cancer), and 6 as category 5 (very high probability of clinically significant cancer).

Within the group of patients with a suspicious lesion (PI-RADS categories 3, 4, and 5), analysis of variance (ANOVA) was used to test for correlation of ADC_4_ measurements between the three categories. Statistically significant differences were observed between groups 3 and 4 and between groups 4 and 5 (p = 0.001), showing a direct correlation between ADC and tumor aggressiveness as classified by PI-RADS™. In ADC_12_ measurements, similar correlations were observed between groups 3 and 4 and between groups 3 and 5.

Of the 158 patients included, 52 underwent prostate biopsy with the following results: 28 (53.8%) with confirmed cancer, 14 (27.0%) negative for cancer, 7 (13.4%) diagnosed with prostatitis, 2 (3.8%) with atypical small acinar proliferation (ASAP), and 1 (2.0%) with PIN (prostatic intraepithelial neoplasia).

The correlation between Gleason score and ADC_4_ and ADC_12_ values was calculated and presented in Table 3. Gleason scores were pooled to facilitate analysis: score 7 included both the results 3 + 4 and 4 + 3, while score 9 included both the results 4 + 5 and 5 + 4.

**Table 3.** Correlation between Gleason score and ADC values

| ADC | Gleason 6 | Gleason 7 | Gleason 8 | Gleason 9 |
| --- | --- | --- | --- | --- |
| Mean ADC<sub>4</sub> | 1121.6 | 1032.8 | 885.5 | 667.7 |
| Mean ADC<sub>12</sub> | 706.4 | 663.3 | 555.8 | 471.0 |

ADC, apparent diffusion coefficient.

Mean ADC values fell as Gleason score increased, confirming a trend for lower ADC values with increasing pathological aggressiveness. However, in both groups, ANOVA demonstrated no statistically significant differences between Gleason grades in areas classified as lesions.

On analysis of the predictive value of the diffusion sequences for detecting cancer in the prostate as confirmed by histopathology (through ROC curves and subsequent calculation of AUC), both techniques were significantly predictive for cancer; the ADC_4_ sequence had a minimum cut-off value of 1153 × 10^−6^mm^2^/s, sensitivity of 71.4% and specificity of 72.3%, and the ADC_12_ sequence, a minimum cut-off value of 637.5 × 10-6 mm^2^/s, sensitivity of 53.6%, and specificity of 82.6% (Figure 6).

**Fig. 6.**
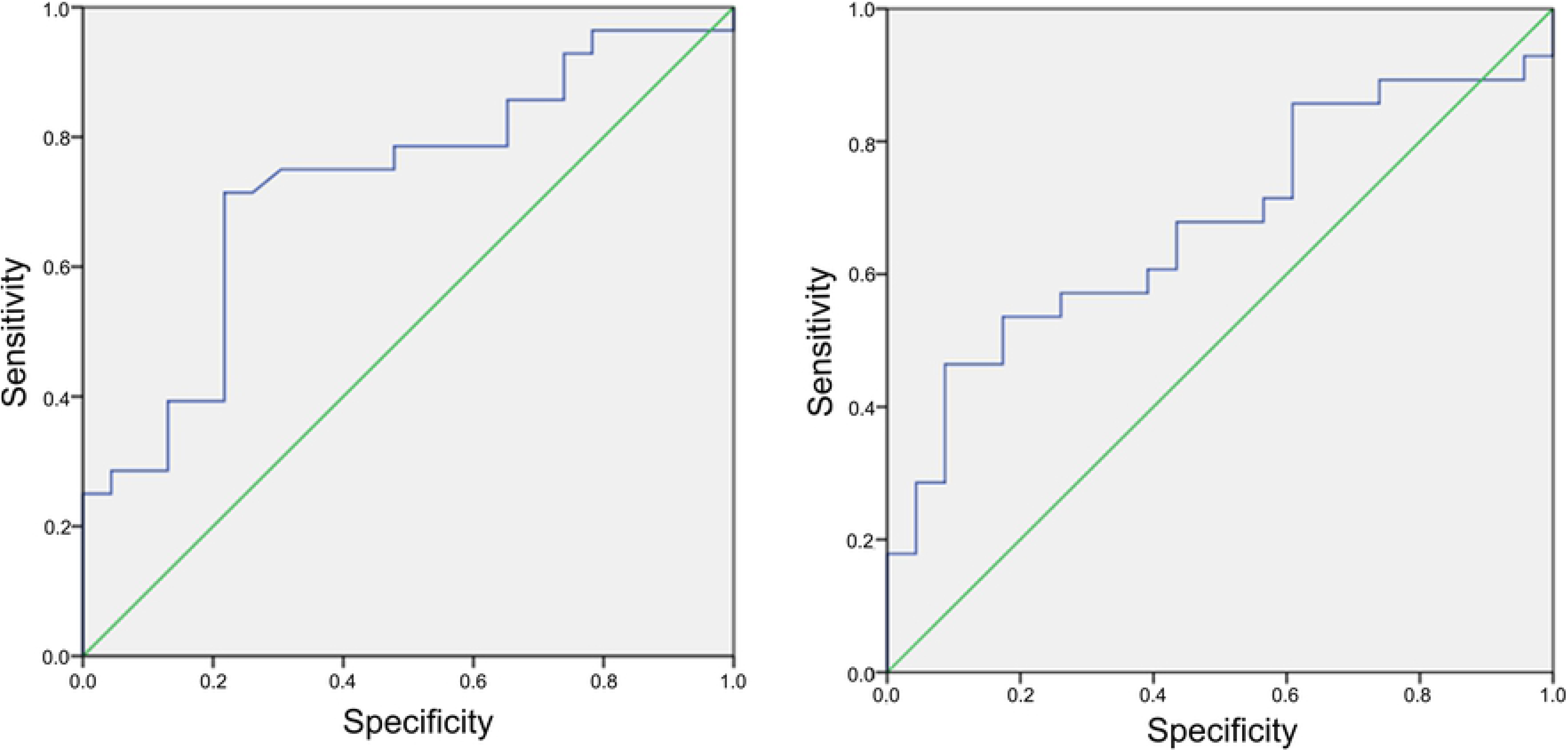
ROC curves for ADC_4_ (right) and ADC_12_ (left). ADC, apparent diffusion coefficient.

The analysis of interobserver agreement related to the classification of image quality and sharpness were made by the kappa coefficient and Spearman correlations, that revealed low but significant agreement across all parameters, except for correlation of the ADC_12_ anatomy classification, which did not demonstrate agreement that was significantly different from zero. In short, the two observers tended to make similar classifications.

On comparative analysis between ADC_4_ and ADC_12_ in relation to anatomy and lesion identification, we obtained significantly higher mean classification values for both observers with ADC_12_ than with ADC_4_ (p < 0.001) (Figure 7), demonstrating that the new technique provides a higher degree of sharpness than the standard sequence, both to study the anatomy of the prostate and to identify suspicious lesions (Figure 8).

**Fig. 7.**
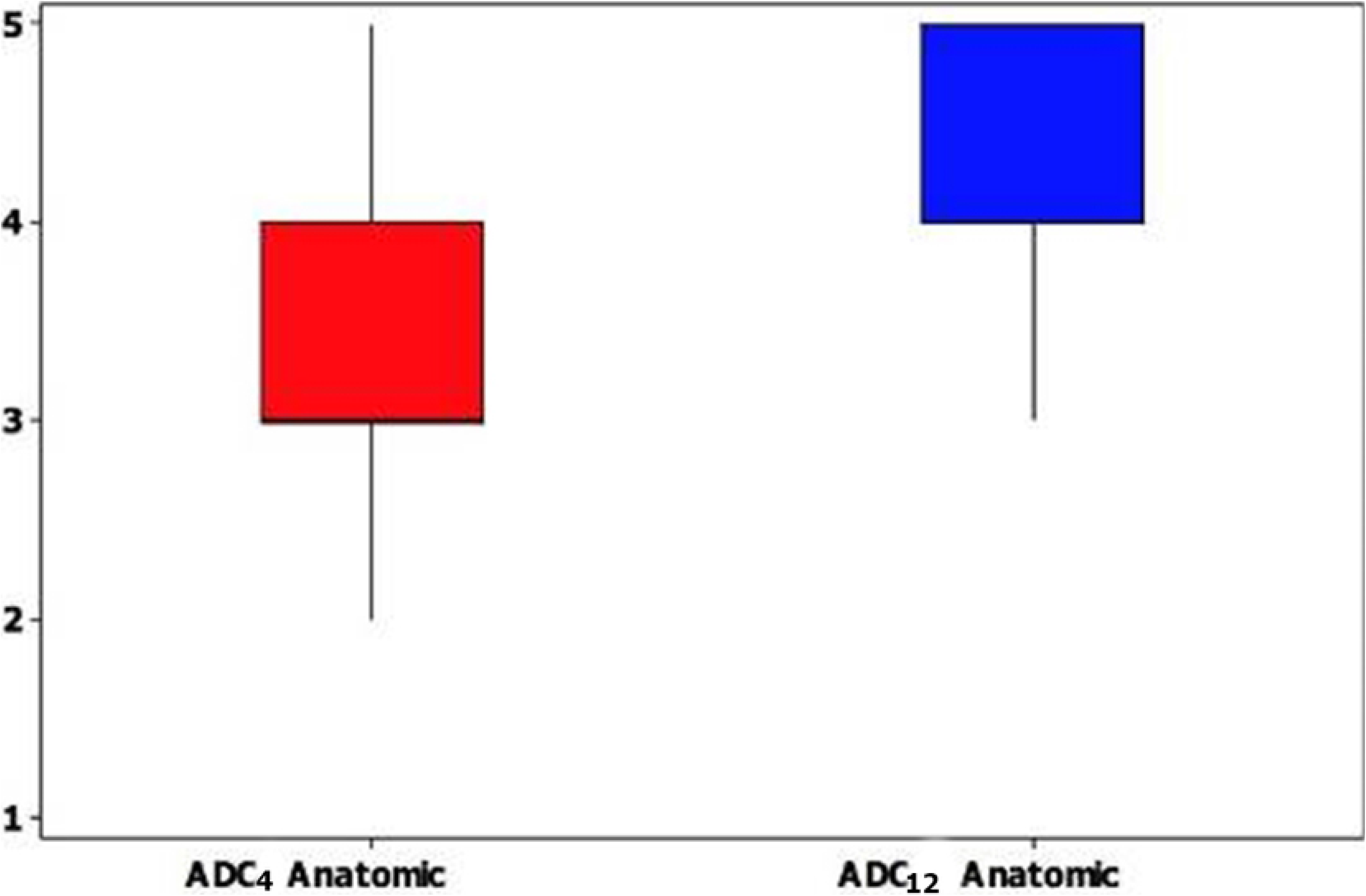
Box plot of Anatomic ADC_4_ and Anatomic ADC_12_. ADC, apparent diffusion coefficient.

**Fig. 8.**
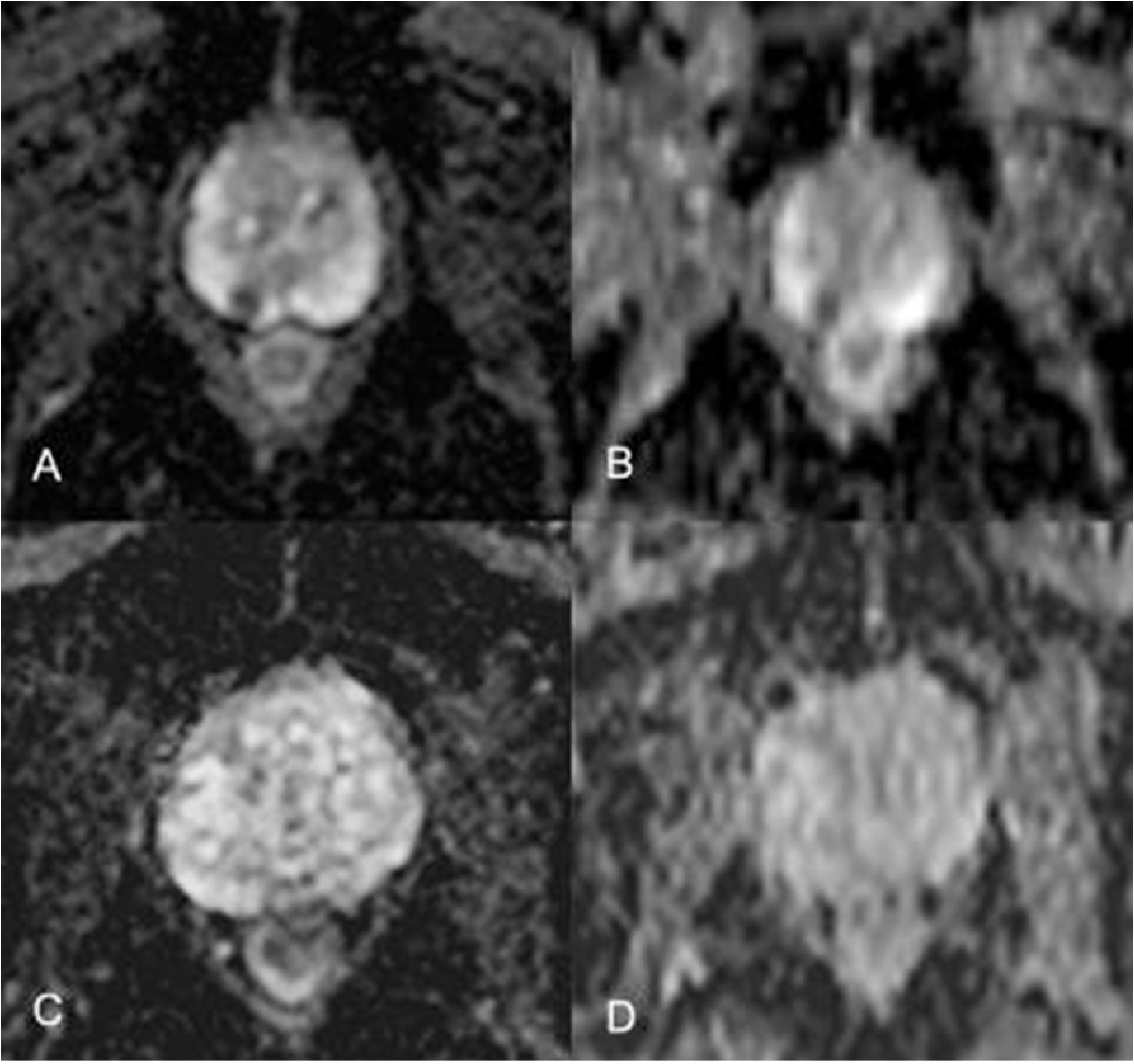
Comparison of image quality between the two diffusion techniques. Panels A and C show characterization of a focal lesion and prostate anatomy, respectively, with an ADC_12_ sequence; panels B and D show the same lesion and anatomy imaged with an ADC_4_ sequence. ADC, apparent diffusion coefficient.

## Discussion

Diffusion imaging is already established in literature and practice as an important tool for study of the prostate, able to function even as a biomarker of tumor aggressiveness [12–17]. The present study demonstrated the viability of a diffusion sequence with 12 *b* values as compared to the standard diffusion sequence of 4 *b* values for routine mp-MRI of the prostate.

Diffusion sequences with several numbers of *b* values have been reported in the recent literature [19–22], but there is no established consensus as to the technical parameters of choice for the study of the prostate. The latest version of PI-RADS™ (2015) does not provide any indication of how many *b* values should be used. Ultra-high *b* values may add some benefit in identification of lesions, but at the expense of decreased image quality and sharpness [20, 21]. In addition, other studies on diffusion sequences of the prostate have described an overlap between benign prostatic focal lesions that restrict diffusion, such as nodules of benign hyperplasia and focal areas of prostatitis, and malignant lesions [15–17]. In the literature review, no studies were found that compared two diffusion sequences with different *b* values in relation to image quality.

In this study, the main objective was to improve the technical parameters and image quality of diffusion sequencing while maintaining its high diagnostic value.

As the primary result, the conspicuity and sharpness of images obtained by diffusion with 12 *b* values were significantly greater than those of images obtained with four *b* values, both for evaluation of normal anatomy and of focal lesions. Thus, interpretation of these images for characterization of possible suspicious lesions was significantly improved, overcoming a challenge that is frequently reported in the literature. This improvement in image quality, with little impact on sequence duration (about 10 additional seconds), greatly optimizes mp-MRI protocol. As the literature increasingly tends to favor MRI for initial screening of prostate cancer, consequently increasing the importance of T2 and DWI sequences [23, 24], this optimization will be of great value to make MRI more effective in identifying lesions.

Both in the standard technique and in the new technique, overall ADC values were statistically significantly lower in neoplastic tissue compared to normal prostate tissue. In addition, ADC values in the 12 *b*-value sequence were comparable to those obtained with four *b* values and were similarly distributed among patients, thus yielding a constant ratio of ADC values in both techniques. On PI-RADS™ classification correlation, both techniques were effective in differentiating between classifications of lower aggressiveness (3) and greater aggressiveness (4 and 5).

On analysis of predictive value, both sequences proved to be significant predictors of cancer, with ADC_12_ having a higher specificity than ADC_4_, demonstrating that it is the technique best able to rule out the possibility of cancer.

Although a statistically significant correlation between ADC_4_ or ADC_12_ values and Gleason score obtained through biopsy was not found, probably due to the small number of patients in our sample who underwent histopathological study, average absolute ADC values decreased as Gleason score increased, demonstrating a trend for mean ADC values to follow the grade of pathological aggressiveness of the tumor.

Major limitations of this study include the technique of histopathological study, which was performed on specimens obtained by cognitive fusion biopsy. Classically, the gold-standard method of pathological study is prostatectomy. Thus, Gleason classifications could have been different in some cases, which might have led to a different percentage of more or less aggressive tumors. Another modality that is currently being studied and which has shown promising results is prostate biopsy with cognitive fusion of MRI and ultrasound imaging, which aids in adequate localization of the suspected lesion and allows identification of additional tissue fragments from the area that is most abnormal on MRI [7, 8, 25]. Another limitation was the small number of patients whose biopsy was positive for prostate neoplasm (n=28), among whom only 17 had a Gleason score of 7 or higher, significantly reducing the number of cases of clinically significant cancer according to the PI-RADS™ criteria. A greater number of cases with proven cancer would be needed to establish statistically relevant correlations with Gleason classification.

## Conclusions

To conclude, the ADC sequences with 12 *b* values are fully viable for MRI of the prostate and produce images of superior quality and sharpness than current techniques, which usually employ four *b* values. Its constant relationship with the standard diffusion sequence and good correlation with the PI-RADS™ and Gleason classifications endorse its use in mp-MRI of the prostate.

## Notes

Conflict of Interest: The authors have no conflicts of interest to disclose.

